# Sex-Specific Neural Adaptations to Acute and Chronic Restraint Stress in Mice

**DOI:** 10.1101/2025.05.20.655203

**Authors:** Ai-Jun Li, Megan McGraw, Hailey Landsparger, Lauren Benjamin, Emily Qualls-Creekmore

**Affiliations:** Department of Integrative Physiology and Neuroscience, Washington State University, Pullman, Washington, USA

**Keywords:** restraint stress, sex differences, c-Fos, desensitization, brain

## Abstract

Stress responses are essential for coping with immediate threats and maintaining physiological homeostasis. While acute stress activates adaptive neuroendocrine and behavioral mechanisms, chronic stress leads to desensitization of these responses, disrupting hormone secretion, neuronal activity, and behavior. Chronic stress is a well-established risk factor for neuropsychiatric disorders, many of which show distinct prevalence and presentation patterns between sexes. However, the neurobiological mechanisms underlying these sex-dependent effects remain poorly understood. This study investigated how acute and chronic stress differentially affect neural activation patterns in male and female mice, with the hypothesis that sex-specific adaptations to chronic stress underlie divergent vulnerabilities to neuropsychiatric disorders. We employed three experimental groups: a control group (no stress), an acute stress group (one hour of restraint stress), and a chronic stress group (one hour of restraint stress daily for ten days). Neural activity was assessed by quantifying c-Fos-positive cells using immunohistochemistry. Acute stress induced widespread neural activation in both sexes, with notable sex differences in c-Fos expression in regions of the hypothalamus, amygdala, and midbrain. Chronic stress led to the desensitization of neuronal activity in most of these regions. Notably, chronically stressed females exhibited more desensitization in specific hypothalamic and amygdaloid regions compared to males. Despite this, corticosterone release remained elevated in stressed females, indicating a decoupling of hormonal and neural responses. These findings suggest that chronic stress elicits distinct neural adaptations in males and females, potentially contributing to the sex-specific vulnerability to neuropsychiatric disorders. Understanding these mechanisms may inform targeted interventions for stress-related pathologies.

## Introduction

Acute physical or psychological stress triggers a series of reactions to counteract immediate threats, cope with the stressor, and maintain body homeostasis^1–4^. These stress responses include sympatho-adrenomedullary and hypothalamic-pituitary-adrenal responses. Studies from both clinical settings^5–8^ and experimental animal models^9–13^ have demonstrated sex differences in these stress responses to an acute stressor. In addition, repeated or chronic stress can result in desensitization of these stress responses, from neuronal activity in the brain to hormone release^14–18^, and desensitization can also present in a sex-dependent manner^19, 20^. One of the main characteristics of stress-induced desensitization is that neuronal activity in stress-sensitive brain regions decreases in reactivity to a stress stimulus that has been experienced repeatedly. The brain is an important target of stress, as it plays a critical role in sensing, relaying, and integrating stress signals to facilitate the systemic stress response. However, the mechanisms by which different brain regions coordinate to adapt to stress, and how these processes become desensitized during chronic stress, remain poorly understood, particularly regarding sex-dependent differences in neuronal activity in response to acute and chronic stress.

Chronic stress has a significant negative impact on the body, both physically and psychologically. Indeed, chronic or repeated physical or psychological stress is known to contribute to the development of neuropsychiatric disorders, such as anxiety, depression, post-traumatic stress disorder (PTSD), and schizophrenia in humans^2, 4, 21, 22^. Moreover, chronic stress can also lead to systemic diseases outside the central nervous system (CNS), such as cardiovascular diseases, gastrointestinal diseases, and metabolic dysfunction^23–26^. In addition, the prevalence of neuropsychiatric disorders is markedly different in men and women. Men are at increased risk for attention deficit hyperactivity disorder and schizophrenia^27, 28^. While depression and PTSD are twofold more prevalent in women^29, 30^. Data from animal studies have also shown that chronic psychosocial and restraint stress cause anxiety-like behaviors, depression-like behaviors, and cognitive impairment in rodents^31–34^, with sex-dependent results^35^. Despite these well-known sex-dependent differences in responses to acute and chronic stress in animals and humans, the underlying mechanisms of these sexual dimorphisms are still unclear.

An expanding body of literature has highlighted sex differences in stress responses across molecular, hormonal, and behavioral domains. At the molecular level, males and females exhibit differential expression of stress-related genes, including glucocorticoid receptors and neuropeptides such as corticotropin-releasing hormone (CRH)^36^. Hormonally, the regulation and reactivity of the hypothalamic-pituitary-adrenal (HPA) axis differ between sexes, with females often showing heightened and prolonged glucocorticoid responses to stressors^37^. Behaviorally, these molecular and endocrine differences translate into distinct coping strategies, susceptibility profiles, and behavioral outcomes following stress exposure^38^. Despite growing recognition of these sex-specific features, the underlying neural activity patterns that support them remain poorly characterized, particularly under conditions of chronic stress, which engages distinct neuroadaptive processes compared to acute stress.

Brain regions such as the hypothalamus, amygdala, midbrain, and hindbrain are central to the coordination of stress responses^39^. The hypothalamus orchestrates HPA axis activation, the amygdala mediates emotional and threat-related processing, and midbrain and hindbrain structures regulate autonomic and behavioral components of the stress response^40^. However, the dynamic interplay among these regions under repeated stress exposure, and how these patterns may diverge between sexes, remains insufficiently understood. This gap limits our ability to pinpoint the neurobiological substrates that contribute to sex-specific resilience or vulnerability to stress-related disorders. To address this knowledge gap, immediate early genes such as c-Fos offer a valuable tool for mapping neuronal activation in response to stress. As a transcription factor rapidly upregulated by synaptic stimulation, c-Fos serves as a reliable proxy for neuronal activation in defined time windows^41^. Quantitative mapping of c-Fos-positive cells across brain regions allows researchers to infer which circuits are recruited by acute versus chronic stress and to identify sex differences in regional activation patterns. Moreover, patterns of co-activation and alterations in connectivity between regions can reveal reorganization of functional networks, a hallmark of chronic stress-induced plasticity. Investigating these sex-dependent changes in both localized activity and network-level integration is essential for a mechanistic understanding of how chronic stress differentially shapes brain function in males and females.

The present study aimed to characterize how acute and chronic stress affect neural activation patterns in male and female mice. We hypothesized that chronic stress induces sex-specific adaptations in neural activity and brain connectivity, contributing to differential vulnerability to neuropsychiatric disorders. To test this, we exposed mice to either a single session of restraint stress (acute stress) or repeated daily restraint over ten days (chronic stress), and quantified neuronal activation via c-Fos immunohistochemistry. We also assessed corticosterone levels to examine hormonal responses and investigated alterations in communication between key brain regions. Through this approach, we sought to elucidate the neural substrates underlying sex-dependent stress responses and provide insight into the mechanisms driving sex biases in stress-related pathologies.

## Methods

### Animals

Male and female C57BL/6J mice aged 8- to 9-weeks-old at the beginning of experiments from Jackson Laboratories (Bar Harbor, ME) were used. Mice were housed at 21- 22° C on a 12-hour light/dark cycle (lights on at 07:00 AM), two mice per cage. Food and water were available ad libitum except during stress sessions. All experimental procedures conformed to National Institutes of Health guidelines and were approved by the Washington State University Institutional Animal Care and Use Committee.

### Restraint stress

Mice in the acute (1 session on day 10) or chronic (daily sessions for 10 consecutive days) restraint stress group were immobilized in a plastic decapicone bag (Braintree Scientific Inc., Braintree, MA) for one hour without access to food or water, as described in previous publications for mice and rats^42, 43^. All restraint stress was performed between 9:00 AM to 11:00 AM (zeitgeber time = ZT2 - ZT4). Body weight (BW) was measured daily before the stress sessions. Mice in the control group did not undergo restraint but were handled daily during BW measurement.

### Tissue collection and Immunohistochemistry

Under deep isoflurane-induced anesthesia, 90 min after the end of the last restraint stress (or handling of the control mice), mice were perfused with ∼15 mL blood bank saline (Fisher Scientific, Hanover Park, IL), followed with ∼15 mL 10 % neutral buffered formalin solution (Sigma-Aldrich Inc., St. Louis, MO). Brains were removed and postfixed in 10 % neutral buffered formalin overnight at 4° C, then cryoprotected in 30 % sucrose in PBS for 24 hours at room temperature. Brains were sectioned coronally into four serial sets (30 µm thickness). Sections were stained using standard avidin-biotin-peroxidase techniques for detecting c-Fos expression^44, 45^. Sections were first incubated in rabbit anti-c-Fos antibody (1:500; MilliporeSigma, Burlington, MA), diluted 1:1000 in 3 % normal donkey serum (NDS) in PBT (0.25 % Triton X-100 in PBS) for 2 days at 4° C. After washes, an overnight incubation in biotin-conjugated donkey anti-rabbit IgG (1:500; Jackson ImmunoResearch) in PBT, and more washes, nickel-intensified diaminobenzidine was used in a peroxidase reaction to produce a gray/black reaction product for c-Fos. After washing, sections were then mounted on SuperFrost slides (VWR, Radnor, PA) and cover slipped for microscopic evaluation. All antibodies used in the experiment were titrated prior to use to determine optimal concentrations.

### Quantification of c-Fos-positive cells

c-Fos-positive cells were counted in the brain regions described below. Brain regions are defined as in *The Mouse Brain in Stereotaxic Coordinates*^46^. Images of c-Fos staining were taken for each region from each mouse using a Zeiss Axio 2 microscope (Carl Zeiss Microscopy GmbH; Oberkochen, Germany). c-Fos-positive cells were counted bilaterally in two coronal sections for each region per mouse, using ImageJ software (Fiji, NIH). The two sections for each region were chosen between the following bregma distance according to the above atlas (distance in mm from bregma, + or – indicates rostral or caudal to bregma, respectively): lateral septum (LS; +0.74 mm to +0.50 mm; including ventral, intermediate and dorsal parts); bed nucleus of stria terminalis (BNST; +0.26 mm to +0.14 mm including medial and lateral division); paraventricular nucleus of hypothalamus (PVH; -0.82 mm to –0.94 mm); lateral hypothalamic area (LHA; -1.22 mm to –1.46 mm); dorsomedial hypothalamic nucleus (DMH; -1.82 mm to –2.06 mm); arcuate nucleus of hypothalamus (Arc; -1.58 mm to –1.82 mm); basolateral amygdala (BLA; -1.22 mm to –1.58 mm); central amygdala (CeA; -1.22 mm to –1.58 mm); basomedial amygdala nucleus (BMA; -1.22 mm to –1.58 mm); medial amygdala (MeA; -1.22 mm to –1.58 mm); paraventricular nucleus of thalamus (PVT; -1.70 mm to –1.94 mm); lateral habenular nucleus (LHb; -1.70 mm to –1.94 mm); ventral tegmental area (VTA; -3.08 mm to -3.16 mm); Locus coeruleus (LC; -5.34 mm to –5.52 mm); nucleus of solitary tract (NTS; -7.48 mm to –7.64 mm).

#### ELISA

Just before the perfusion as in the above section, blood samples were collected from the right ventricle of heart, with a 26G needle and 1 mL syringe. After incubation on ice for 1 hour, serum was collected by a centrifuge at 2000g for 15 min at 4° C and then stored at -80° C. Corticosterone levels were measured with an ELISA assay (Enzo Life Sciences Inc., Farmingdale, NY), according to the manufacturer’s protocol and as previously described^45^. The intra- and inter-assay coefficient variation (CVs) for corticosterone assay were 6.6, 8.4, 8.0 % and 7.8, 8.2, 13.1 % for high, medium, low concentrations, respectively, according to the data sheet.

### Network Connections

Inter-regional correlations of c-Fos-positive cell expression was calculated using Spearman’s correlation matrices for male and female mice undergoing control, acute, and chronic stress conditions. To assess potential network connectivity, *r*-values underwent Fisher’s r-to-z transformation to convert correlation values to comparable *z*-scores. The difference and significance between the *z*-scored correlation coefficients was calculated between male and female chronic and acute stress conditions. This was used to assess changes in network connections between stress conditions.

### Statistical Analysis

All results are presented as mean ± SEM. One-way ANOVA, two-way ANOVA, three-way repeated measures ANOVA, and Two-tailed z-tests were used for statistical analysis. After significance via one-two-three-way ANOVA was determined, post hoc multiple comparisons of individual groups were used (Tukey multiple comparisons test, or Fishers LSD depending on number of groups). A *p*-value < 0.05 was considered statistically significant, and notated as follows: * p < 0.05, ** p < 0.01, *** p < 0.001, unless otherwise stated.

## Results

### c-Fos expression was measured in 15 brain regions following restraint stress

The timeline of the experiments is shown in **Fig. 1A**, wherein mice received ten consecutive days of daily handling (DH) or one hour of restraint stress (RS) and body weight measurement. On the last day, 90 min after DH or RS, blood samples and brain tissues were collected from the control, acute, and chronic stress groups, both males and females. Corticosterone levels were measured with ELISA and c-Fos-positive cells were counted in the brain regions as shown in **Fig. 1B**.

**Figure 1.**
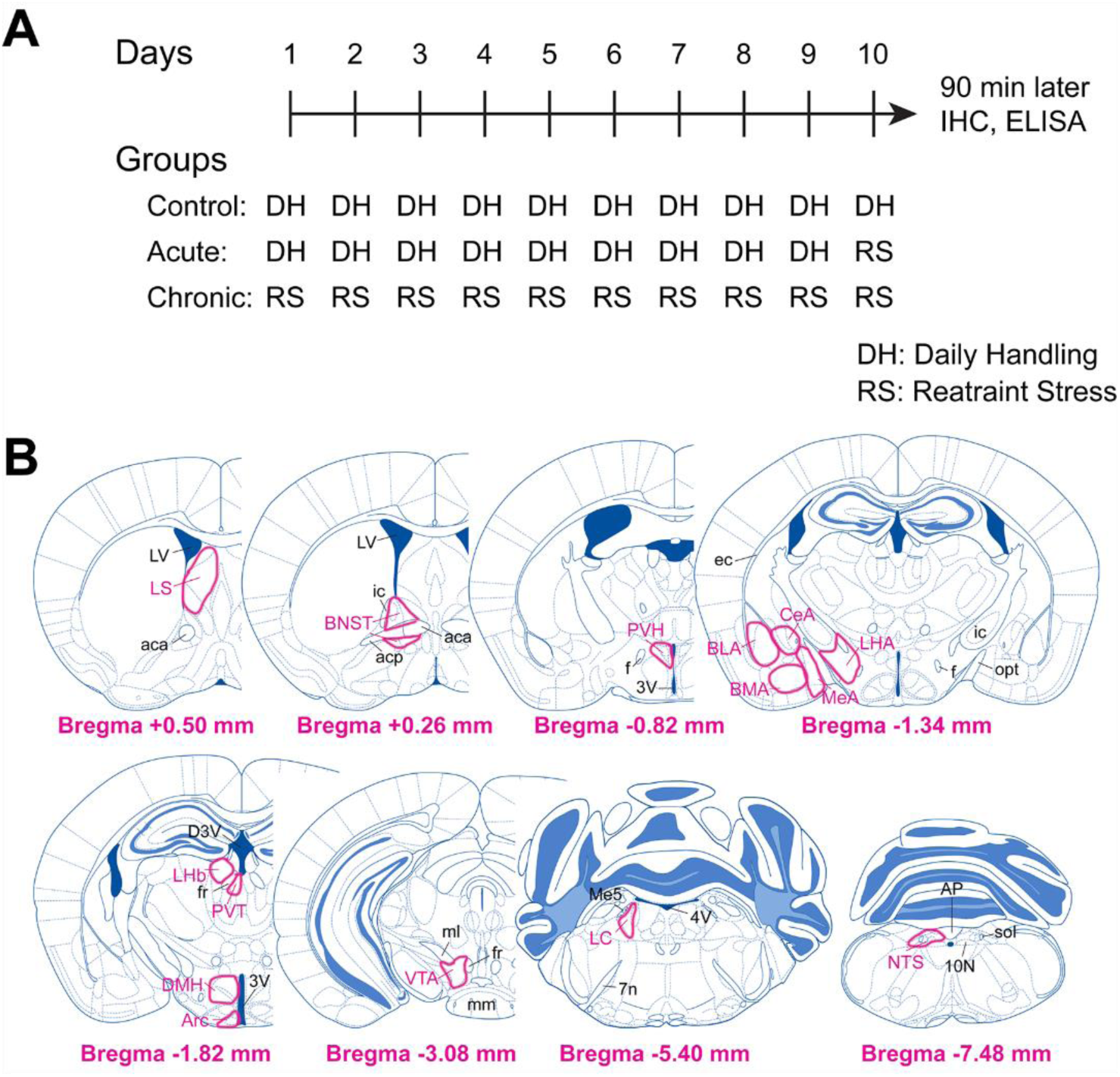
Timeline of experiments and mice brain regions for c-Fos counting. **A.** Timeline of experiments for the control, acute, and chronic stress groups. Both male and female mice were handled or stressed (one hour restraint) daily for ten consecutive days. Mice were perfused 90 min after the last handling or restraint stress session. Expression of c-Fos in brain regions was counted by immunohistochemistry (IHC) methods and corticosterone levels in the blood samples were detected by ELISA. **B.** brain regions where c-Fos-positive cells were counted. Distance rostral (+), or caudal (-) to Bregma is shown. 3V, third ventricle; 4V, fourth ventricle; 7n, facial nerve; 10N, dorsal motor nucleus of vagus; aca, anterior commissure, anterior part; acp, anterior commissure, posterior part; AP, area postrema; Arc, arcuate nucleus of hypothalamus; BLA, basolateral amygdala; BMA, basomedial amygdala; BNST, bed nucleus of stria terminalis; CeA, central amygdala; D3V, dorsal 3V; DMH, dorsomedial hypothalamic nucleus; ec, external capsule; f, fornix; fr: fasciculus retroflexus; ic, internal capsule; LC, locus coeruleus; LHA, lateral hypothalamic area; LHb, lateral habenular nucleus; LS, lateral septum; LV, lateral ventricle; Me5, mesencephalic trigeminal nucleus; MeA, medial amygdala; ml, medial lemniscus; mm, medial mammillary nucleus; NTS, nucleus of solitary tract; opt, optic tract; PVH, paraventricular nucleus of hypothalamus; PVT, paraventricular nucleus of thalamus; sol, solitary tract; VTA, ventral tegmental area.

### Chronic stress decreases body weight in both males and females

One hour daily repeated restraint stress (chronic stress) decreased the body weight (BW) in male and female mice (**Fig. 2A**, Three-way repeated measures ANOVA, N = 8 mice per group, F_(1, 28)_ = 30.27, p < 0.0001). In addition to a main stress effect, there was an interaction of stress and time (**Fig. 2A**, Three-way repeated measures ANOVA, N = 8 mice per group, F_(9, 252)_ = 19.02, p < 0.001), but no effect of sex (**Fig. 2A**, Three-way repeated measures ANOVA, N = 8 mice per group, F_(1, 28)_ =0.001, p = 0.9749) or an interaction between sex and time or treatment. By day 4, the BW gain (%) in the male chronic stress mice group was significantly lower when compared to BW gain (%) within subject prior to onset of chronic stress (**Fig. 2A**, Tukey’s multiple comparisons test, N = 8 per group, T_(280)_ = 3.511), and was significantly lowered compared to control males by day 5 (**Fig. 2A**, Tukey’s multiple comparisons test, N = 8 mice per group, T_(8.03)_ = 4.899, p = 0.0417). In female mice, chronic stress decreased the BW gain (%) compared to baseline by day 8 (**Fig. 2A**, Tukey’s multiple comparisons test, N = 8 per group, T_(280)_ = 4.128, p = 0.0002), and was also decreased compared to control by day 8 (**Fig. 2A**, Tukey’s multiple comparisons test, N = 8 per group, T_(280)_ = 7.31 p = 0.0003).

**Figure 2.**
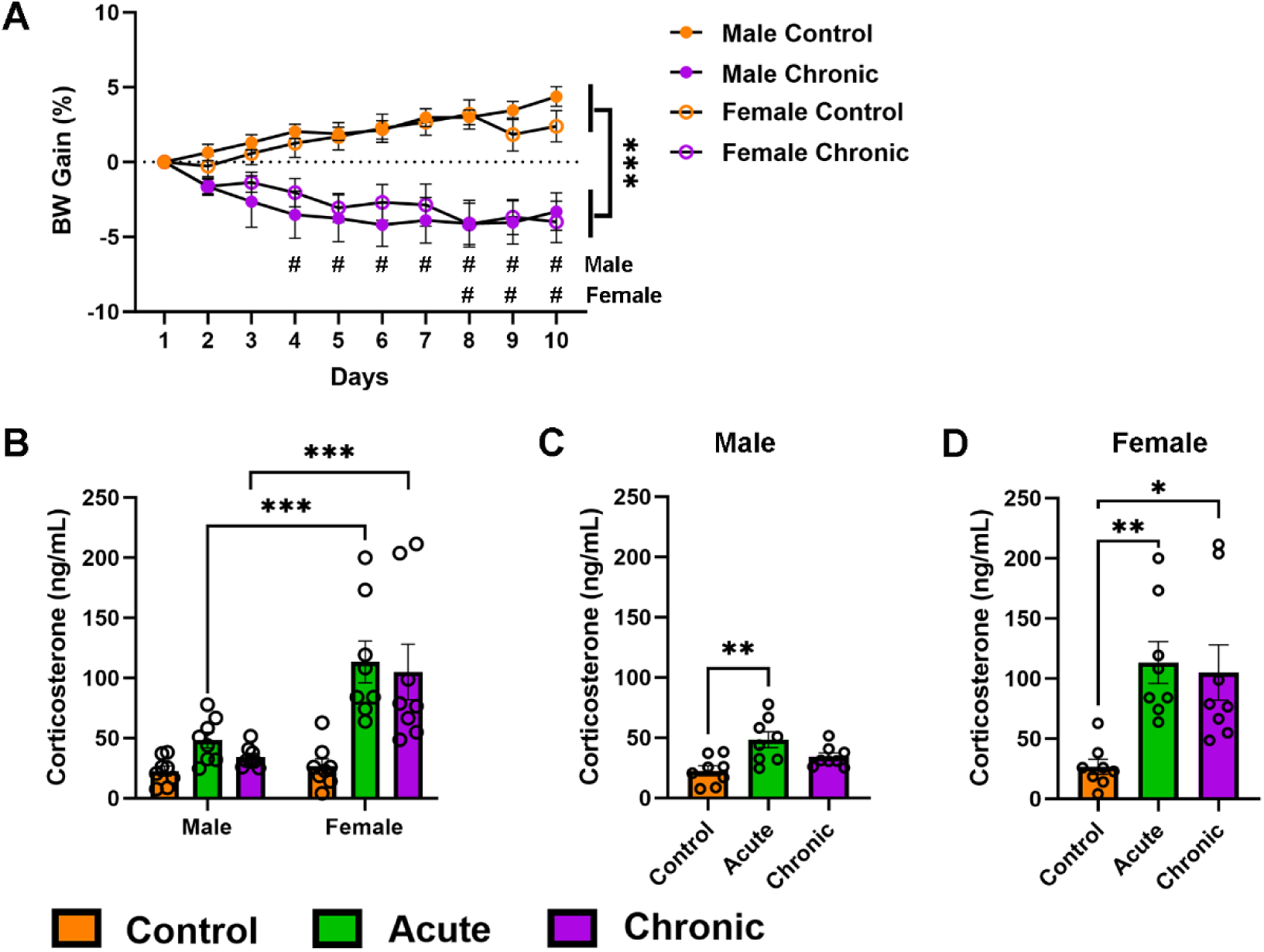
Physiological markers of acute and chronic stress in males and females. **A.** Daily change in body weight (BW gain), reported as percent change, are shown for male and female controls and chronic stress mice. Data are expressed as mean ± SEM, N = 8 mice per group. *** p < 0.0001, main effect of stress; # p < 0.05, Tukey’s multiple comparisons test compared to pre-stress weight; three-way repeated measures ANOVA. **B-D.** Restraint stress increases serum corticosterone levels in males and females. **B.** Female acute and chronic stress CORT levels are significantly increased compared to males, *** p < 0.0001, Tukey’s multiple comparisons test, two-way ANOVA. Male **(C)** and female **(D)** CORT levels are increased following acute stress, and female levels remain elevated following chronic stress, * p < 0.05, ** p < 0.1, Tukey’s multiple comparisons test, one-way ANOVA.

### Corticosterone levels were increased acutely with sex specific differences in chronic exposure

Blood corticosterone levels were measured at day 10, 90 min after the last handling or stress session (**Fig. 1A**). Corticosterone levels were significantly elevated in females compared to males (**Fig. 2B**, Two-way ANOVA, N = 8 per group, F_(1, 42)_ = 20.74, p < 0.001). Stress also increased corticosterone levels (**Fig. 2B**, Two-way ANOVA, N = 8 per group, F_(2, 42)_ = 11.28, p = 0.0001) and there was a significant interaction between sex and stress (**Fig. 2B**, Two-way ANOVA, N = 8 per group, F_(2, 42)_ = 4.356, p = 0.0191). Female corticosterone levels were significantly increased compared to males in the acute and chronic stress group (**Fig. 2B**, Tukey’s multiple comparisons test, N = 8 per group, acute: T_(42)_ = 5.187, p = 0.0007; chronic: T_(42)_ = 5.648, p = 0.0003). However, as the increased variability in the females violated the equal variance assumption of the Tukey’s multiple comparisons test, any effects within the male group were overshadowed. Thus, to ensure male mice did have a significant HPA response to restraint stress, in line with previous literature^45^, one-way ANOVAs were used to compare within sex (**Fig. 2C-D)**. This showed that stress significantly increased corticosterone levels in both male (**Fig. 2C**, One-way ANOVA, N = 8 per group, F_(2, 21)_ = 7.196 p = 0.0042) and female (**Fig. 2D**, One-way ANOVA, N = 8 per group, F_(2, 21)_ = 7.865 p = 0.0028) mice. In males, this effect of stress is driven by a significant increase in corticosterone levels in the acute stress group when compared with the same sex control mice (**Fig. 2C**, Tukey’s multiple comparisons test, N = 8 per group, T_(21)_ = 5.356, p = 0.003). The corticosterone levels in chronically stressed males had no significant difference compared to acute (**Fig. 2C**, Tukey’s multiple comparisons test, N = 8 per group, T_(21)_ = 2.945, p = 0.1179) and control (**Fig. 2C**, Tukey’s multiple comparisons test, N = 8 per group, T_(21)_ = 2.412, p = 0.2266) groups. Whereas, the chronically stressed female corticosterone levels were significantly increased compared to controls (**Fig. 2D**, Tukey’s multiple comparisons test, N = 8 per group, T_(21)_ = 4.594, p = 0.0103), but not acute stressed mice (**Fig. 2D**, Tukey’s multiple comparisons test, N = 8 per group, T_(21)_ = 0.4906, p = 0.936).

### Acute stress increased c-Fos in all regions

In this study, c-Fos-positive cells were counted in 15 brain regions (**Fig. 1B**). Regions were chosen based on a significant increase in c-Fos expression in a region following acute restraint stress. To assess changes in c-Fos expression from acute and chronic stress, the raw c-Fos-positive cell count per region was found for all groups and analyzed by two-way ANOVA looking at main effects of sex, stress, and an interaction between sex and stress followed by post hoc Tukey’s multiple comparisons test (Supplemental Table 1). Then, to assess changes in percent change of c-Fos expression from baseline (c-Fos-positive count – average c-Fos-positive count for same sex control / average c-Fos-positive count for same sex controls * 100), two-way ANOVA was performed to identify significant main effects of sex and chronicity of stress, and an interaction between sex and chronicity followed by post hoc Fisher’s LSD test (Supplemental Table 2). Two-way ANOVA analysis of c-Fos-positive cell count (Supplementary Table 1) confirmed that all brain regions had a main effect of stress (p < 0.001). Two-way ANOVA analysis of percent change in c-Fos-positive expression demonstrated a significant effect of stress chronicity of all regions except the VTA and the LC (Supplemental Table 2).

### Chronic stress-induced c-Fos expression patterns differed by region but similarly between sexes

Based on post hoc multiple comparisons (Supplemental Table. 1-2), parts of the forebrain (LS), hypothalamus (PVH, DMH, and Arc), amygdala (CeA, BMA, and MeA), and the thalamus (LHb) had similar patterns between male and female c-Fos-positive cells counts after chronic restraint. These groups were further delineated into whether c-Fos expression after chronic restraint was fully attenuated compared to control (representative images **Fig. 3**, quantification **Fig. 4**), or if it was attenuated compared to acute restraint but remained significantly increased compared to controls (representative images, **Fig. 5** quantification **Fig. 6**). The remaining regions (BNST, LHA, BLA, PVT, and NTS) all had differing patterns between males and females demonstrating different effects of chronic stress on region activation (representative images **Fig. 7**, quantification **Fig. 8**). For the quantification of each region, the raw count of c-Fos-positive cells for all groups were shown (left), as well as the percent change for acute and chronic conditions compared to same sex controls (right).

**Figure 3.**
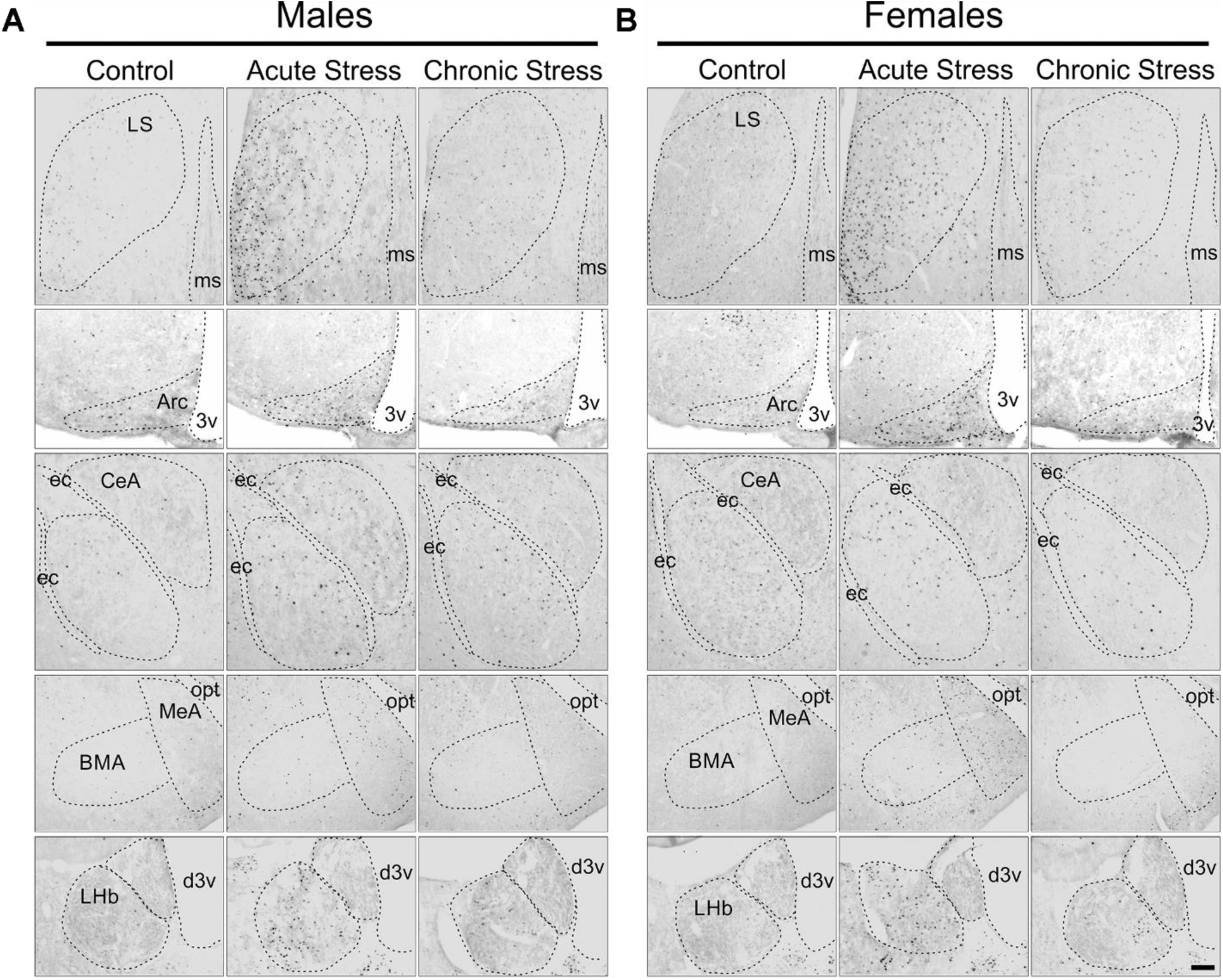
Representative stress-induced c-Fos expression in brain regions exhibiting full attenuation following chronic stress. Representative c-Fos expression in male **(A)** and female **(B)** mice from the control (left), acute (middle), and chronic (right) stress groups. Represented regions all demonstrated similar patterns of attenuation following chronic stress in both sexes. LS, lateral septum; ms, medial septal nucleus; Arc, arcuate nucleus of the hypothalamus; 3v, third ventricle; CeA, central amygdala; ec, external capsule; BMA, basomedial amygdala; MeA, medial amygdala; opt, optic tract; LHb, lateral habenular nucleus; d3v, dorsal third ventricle. Scale bar = 0.1mm for all except for BMA/MeA where the scale bar = 0.16mm.

**Figure 4.**
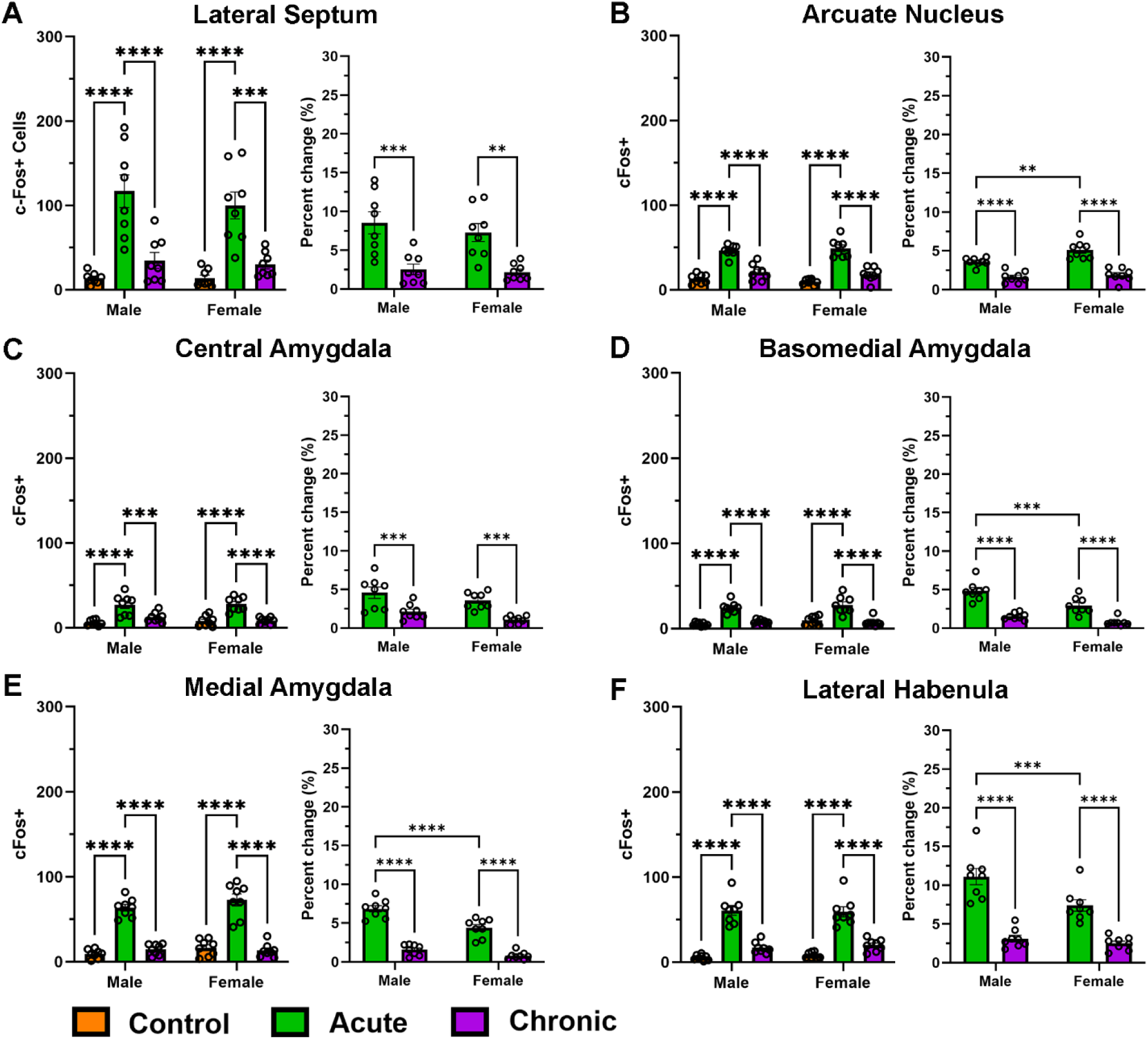
Stress-induced c-Fos expression in male and female brain regions exhibiting full attenuation following chronic stress. c-Fos expression was detected 90 min after the last stress session from the control, acute, and chronic stress groups. c-Fos-positive cells in each region were counted and graphed (left). Percent change of c-Fos expression was then calculated based on the average c-Fos-positive cell count of same sex controls and graphed per region (right). All data is presented as mean ± SEM. N = 8 mice per group. * p < 0.05, ** p < 0.01, *** p < 0.001, Tukey’s multiple comparisons test for c-Fos-positive cell count (left) and Fisher’s LSD test for percent change (right), two-way ANOVA.

**Figure 5.**
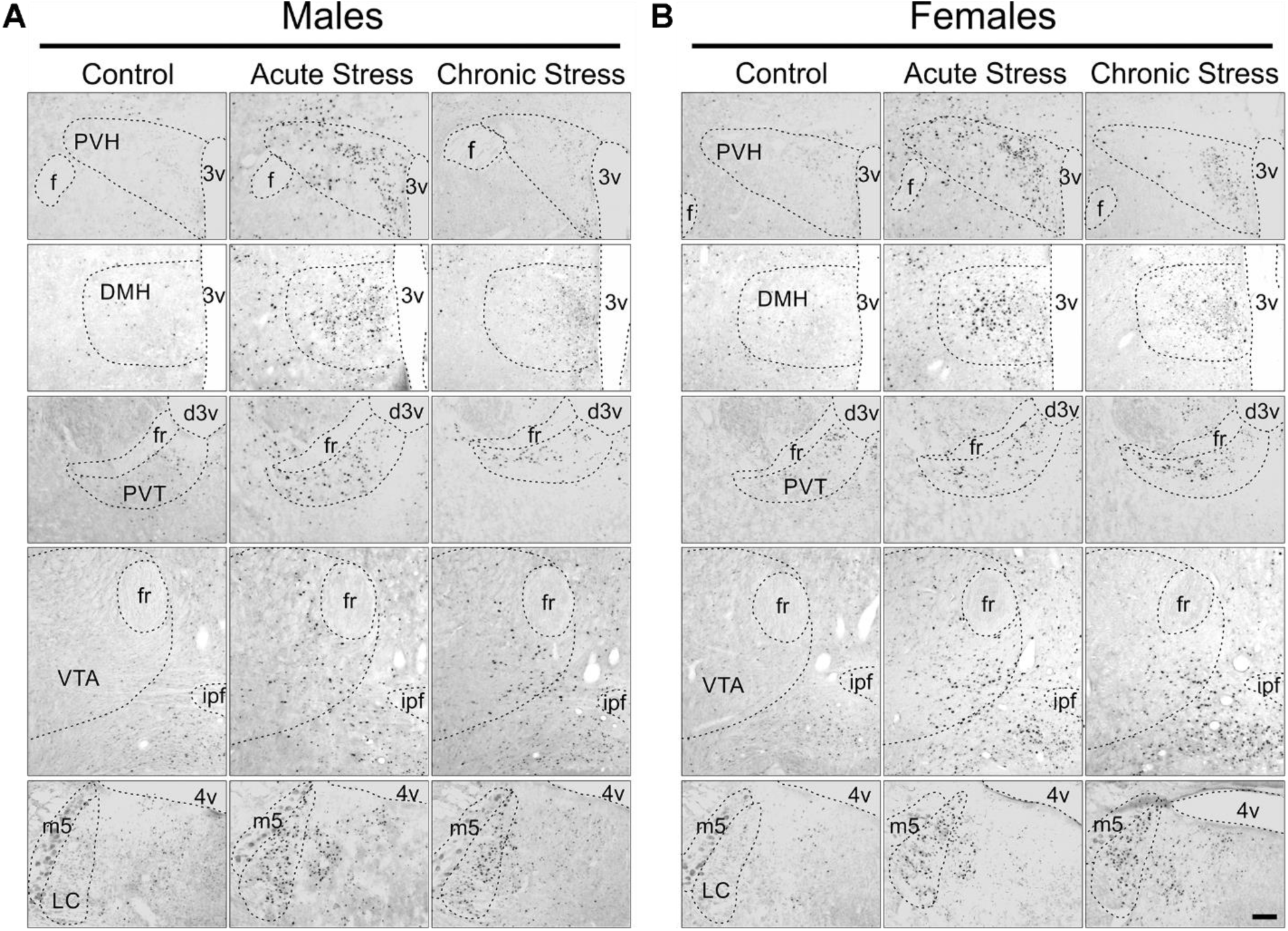
Representative stress-induced c-Fos expression in brain regions exhibiting partial attenuation following chronic stress. Representative c-Fos expression in male **(A)** and female **(B)** mice from the control (left), acute (middle), and chronic (right) stress groups. Represented regions all demonstrated similar patterns of partial attenuation following chronic stress in both sexes. PVH, paraventricular nucleus of the hypothalamus; f, fornix; 3v, third ventricle; DMH, dorsomedial hypothalamic nucleus; PVT, paraventricular nucleus of the thalamus; fr: fasciculus retroflexus; d3v, dorsal third ventricle; VTA, ventral tegmental area; ipf, interpeduncular fossa; LC, locus coeruleus; m5, mesencephalic trigeminal nucleus; 4v, fourth ventricle. Scale bar = 0.1mm.

**Figure 6.**
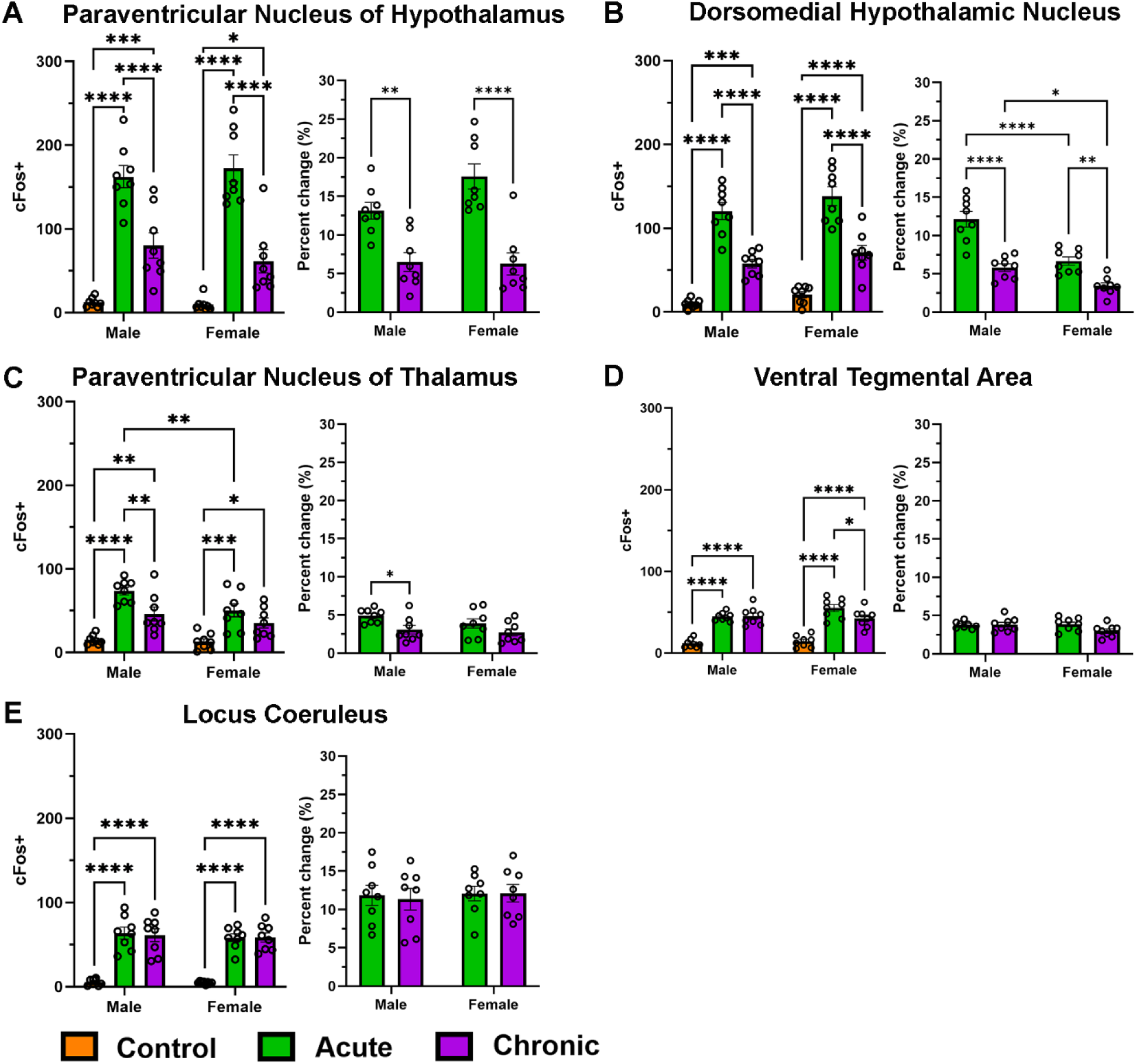
Stress-induced c-Fos expression in male and female brain regions exhibiting partial attenuation following chronic stress. c-Fos expression was detected 90 min after the last stress session from the control, acute, and chronic stress groups. c-Fos-positive cells in each region were counted and graphed (left). Percent change of c-Fos expression was then calculated based on the average c-Fos-positive cell count of same sex controls and graphed per region (right). All data is presented as mean ± SEM. N = 8 mice per group. * p < 0.05, ** p < 0.01, *** p < 0.001, Tukey’s multiple comparisons test for c-Fos-positive cell count (left) and Fisher’s LSD test for percent change (right), two-way ANOVA.

**Figure 7.**
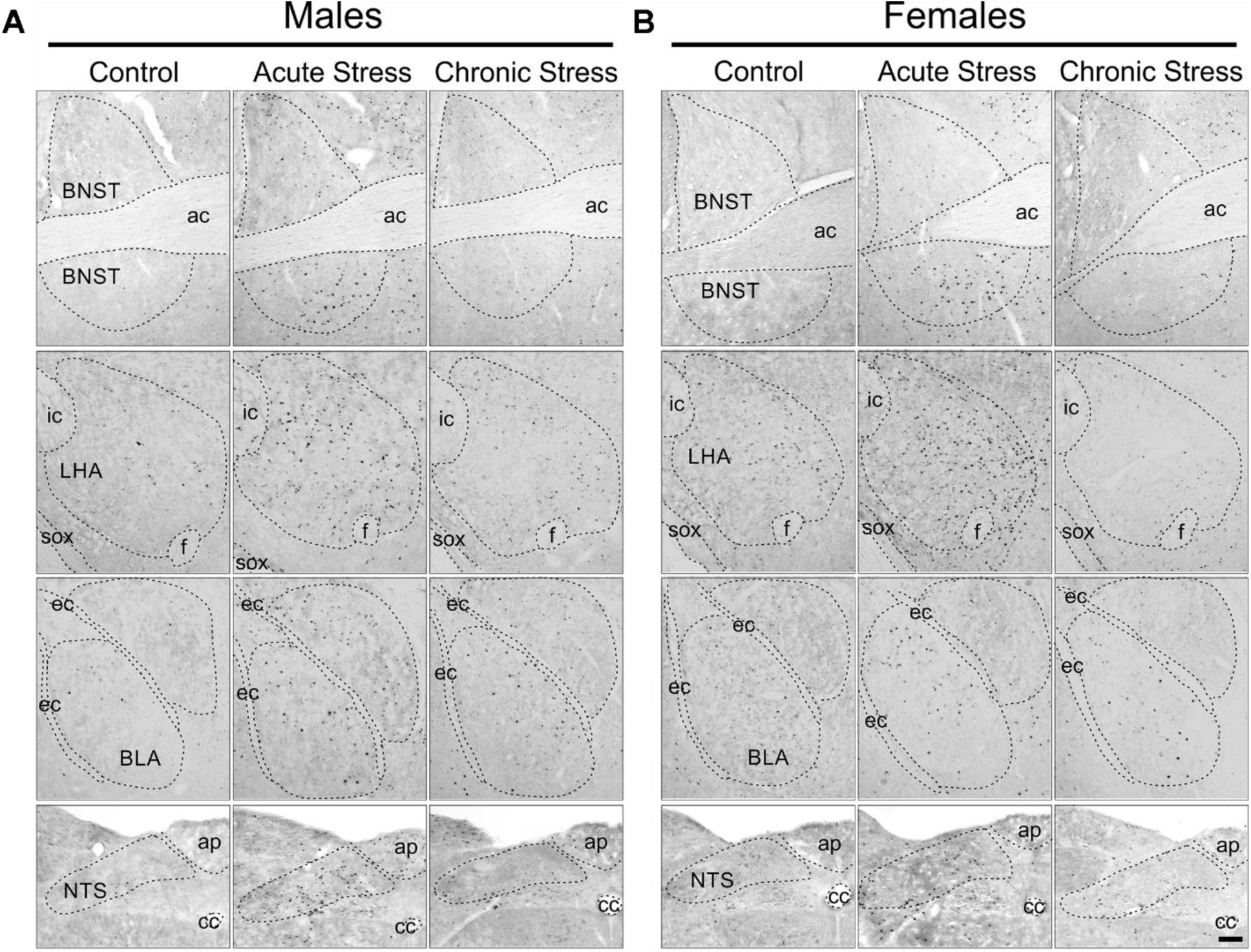
Representative stress-induced c-Fos expression in brain regions exhibiting differing patterns of attenuation between sexes following chronic stress. Representative c-Fos expression in male **(A)** and female **(B)** mice from the control (left), acute (middle), and chronic (right) stress groups. Represented regions demonstrated differing patterns of attenuation following chronic stress between sexes. BNST, bed nucleus of stria terminalis; ac, anterior commissure; LHA, lateral hypothalamic area; ic, internal capsule; f, fornix; sox, supraoptic decussation; BLA, basolateral amygdala; ec, external capsule; NTS, nucleus of solitary tract; ap, area postrema; cc, central canal. Scale bar = 0.1mm.

**Figure 8.**
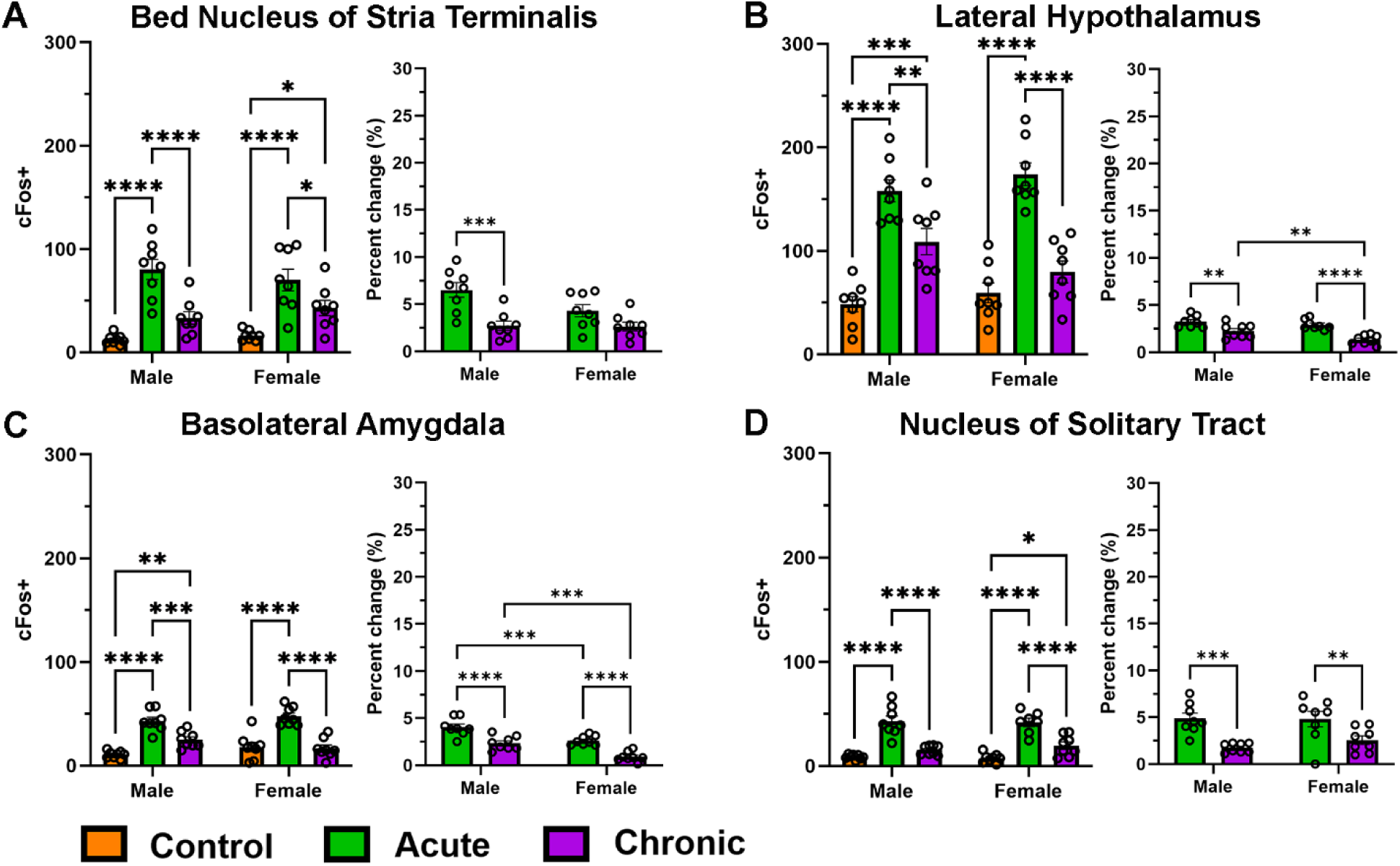
Stress-induced c-Fos expression with differing levels of attenuation between males and females. c-Fos expression was detected 90 min after the last stress session from the control, acute, and chronic stress groups. c-Fos-positive cells in each region were counted and graphed (left). Percent change of c-Fos expression was then calculated based on the average c-Fos-positive cell count of same sex controls and graphed per region (right). All data is presented as mean ± SEM. N = 8 mice per group, except N = 7 for NTS control females. * p < 0.05, ** p < 0.01, *** p < 0.001, Tukey’s multiple comparisons test for c-Fos-positive cell count (left) and Fisher’s LSD test for percent change (right), two-way ANOVA.

Regions that had fully attenuated in both males and females (LS, Arc, CeA, BMA, MeA, and LHb, **Fig. 3**) demonstrated a reliable increase in c-Fos expression acutely in both males and females and a robust decrease in chronic expression compared to acute stress (**Fig. 4**). In all these regions, chronic expression was not significantly different from control levels (Supplemental Table 1). Along with having had similar patterns in males and females in c-Fos-positive cell count, none of these regions had a significant main effect of sex. However, when assessing percent change based on the average c-Fos-positive cell count, with respect to same sex controls, a main effect of sex was found for all regions except for the LS (Supplementary Table 2). In the Arc, this effect was driven by an increase in the percent change of c-Fos expression in the acute group (**Fig. 4B**, Uncorrected Fisher’s LSD, N = 8 per group, T_(28)_ = 3.558, p = 0.0014), while in the CeA (**Fig. 4C**) no one group was significantly different, suggesting there was a difference in cell count not specific to the stress paradigm. On the other hand, the BMA, MeA, and LHb all had a significant decrease in the percent change of c-Fos-positive expression in acutely stressed females compared to males (**Fig 4D-F**). Additionally, the LHb (**Fig. 4F**) and MeA (**Fig. 4E**) had a main interaction between sex and stress (**Fig. 4**, Two-way ANOVA, N = 8 per group LHb: F_(1, 28)_ = 4.941, p = 0.0345; MeA: F_(1, 28)_ = 6.438, p = 0.0170).

Regions that only partially attenuated in both males and females (PVH, DMH, PVT, **Fig. 6A-C**) demonstrated a reliable increase in c-Fos expression acutely in both males and females and a decrease in chronic expression compared to acute. However, compared to control expression, chronic expression remained significantly increased. Additionally, two regions (VTA and LC, **Fig. 6 D-E**) had no attenuation and both acute and chronic expression remained elevated compared to controls, with no difference between stress conditions. While having had similar patterns in males and females in c-Fos-positive cell count, the DMH (**Fig. 6B**) and PVT (**Fig. 6C**) had a significant effect of sex (**Fig. 6**, Two-way ANOVA, N = 8 per group, DMH: F_(1, 42)_ = 4.914, p = 0.0321; PVT: F_(1, 42)_ = 6.160, p = 0.0172). Within the DMH, no one group was significantly different between sexes, but all female groups appeared equally elevated compared to males (**Fig. 6B**). In the PVT, the effect of sex was driven by a decrease in c-Fos-positive cells of acutely stressed females compared to males (**Fig. 6C**). However when comparing percent change in c-Fos expression, the sex effect in the PVH was no longer present (**Fig. 6A**, Two-way ANOVA, N = 8 per group, F_(1, 28)_ = 2.380, p = 0.1341) while the main effects in the DMH were maintained and made more prevalent with a main effect of sex (**Fig. 6B**, Two-way ANOVA, N = 8 per group, F_(1, 28)_ = 34.63, p < 0.0001), and an interaction of sex and stress (**Fig. 6B**, Two-way ANOVA, N = 8 per group, F_(1, 28)_ = 5.223, p = 0.0301). Multiple comparison analysis of percent change in the DMH revealed that, when normalized to control expression, female c-Fos expression was decreased compared to male expression following both acute and chronic stress (**Fig. 6B**, Uncorrected Fisher’s LSD, N = 8 per group, acute: T_(28)_ = 5.777, p < 0.0001, chronic: T_(28)_ = 2.545, p = 0.0167).

### Chronic stress-induced c-Fos expression patterns differed by sex within the same regions

In regions that had differing patterns of attenuation in males and females (BNST, LHA, BLA, and NTS, **Fig. 7**), the BNST (**Fig. 8A**) and NTS (**Fig. 8D**) fully attenuated in the males while the females only partially attenuated; whereas, in the LHA (**Fig. 8B**) and BLA (**Fig. 8C**), the males had partially attenuated while the females had fully attenuated. Interestingly, none of these regions had a main effect of sex when assessing c-Fos-positive cell count (Supplementary Table 1.), but the BLA did have a significant interaction between sex and stress (**Fig. 8C**, Two-way ANOVA, N = 8 per group, F_(2, 42)_ = 3.428, p = 0.0418) demonstrating the divergent attenuation pattern in the BLA. When comparing the percent change, in these regions, the LHA demonstrated a main effect of sex (**Fig. 8B**, Two-way ANOVA, N = 8 per group, F_(1, 28)_ = 5.470, p = 0.0267), that was driven by a decrease in percent change of c-Fos expression in females after chronic stress compared to males (**Fig. 8B**, Uncorrected Fisher’s LSD, T_(28)_ = 2.235, p = 0.0336). When comparing the percent change in the BLA, the only significant main effect was an effect of sex (**Fig. 8C**, Two-way ANOVA, N = 8 per group, F_(1, 28)_ = 33.37, p < 0.0001). Multiple comparisons tests revealed that the sex effect in the BLA was driven by a decrease in c-Fos expression in females compared to males in both the acute and chronic conditions (**Fig. 8C**, Uncorrected Fisher’s LSD, N = 8 per group, acute: T_(28)_ = 4.078, p = 0.0003, chronic: T_(28)_ = 4.091, p = 0.0003).

### c-Fos expression indicates distinct sex-specific network connections that are altered by stress

We further analyzed the correlation coefficient between the c-Fos-positive cell count in each region as an indirect measure of region interconnectedness (**Fig 9**). This was done to determine whether sex-specific stress responsive networks were interrupted by acute or chronic restraint stress. Non-stressed controls demonstrated a stark pattern difference in male and female mice (**Fig. 9A**). Here we saw that males had 5 significant Spearman-r positive correlations between two regions (**Fig. 9A, left**). Females, however, demonstrated 23 significant Spearman-r positive correlations between two regions (**Fig. 9A, right**). When analyzing region interconnectedness during acute stress, in males there were 7 significant positive correlations and 3 negative correlations (**Fig. 9B, left**), with only the positive correlation between the LHA and VTA being conserved. In acutely stressed females, the number of positive correlations was reduced to 5, but no negative correlations emerged (**Fig. 9B, right**). Of these regions, all but the correlation between PVT and LHb were present in control animals, demonstrating that the other 19 correlations were disrupted by acute stress in females. For the animals stressed chronically, males demonstrated 16 positive correlations and only 1 negative correlation (**Fig. 9C left**). Of these correlations, the positive correlations between the LS and CeA as well as the BNST and LHA were maintained, but the connection between the LS and BNST changed from a significant negative correlation to a significant positive correlation. Additionally, the positive correlation between the BLA and CeA was restored. In the chronically stressed females, the number of positive correlations was increased to 8, with no emergence of negatively correlated regions (**Fig. 9C, right**). The interconnection between the LS and PVT as well as the BLA and MeA was maintained throughout all conditions. Additionally, interconnections between the DMH and BLA, BMA, and MeA were restored similarly to control animals, along with the positive correlation between the BLA and VTA.

**Figure 9.**
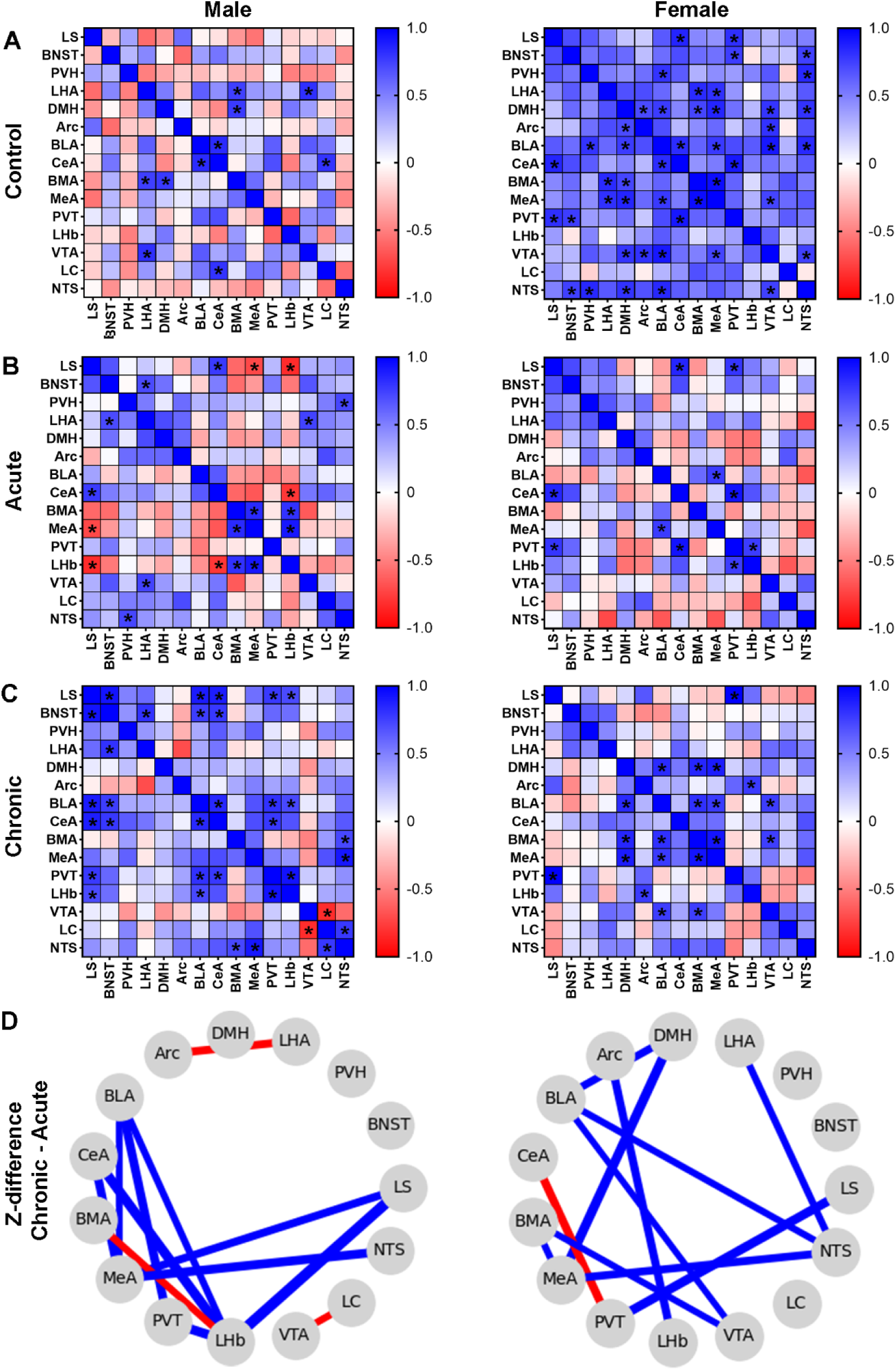
Sex-specific network changes during stress desensitization. Spearman correlation matrices demonstrating inter-regional correlations of c-Fos expression in male (left) and female (right) mice. Correlation coefficients (*r*-values) are represented by blue (positive correlation) and red (negative correlation) colors on a sliding scale of strength (scale, right). Statistically significant inter-regional connections during control **(A)**, acute **(B)**, and chronic **(C)** stress conditions were indicated by * p < 0.05. **D.** The strongest changes in network connectivity (as indicated by the difference in Fisher’s transformed *z*-scores, i.e. *z-*diff, p < 0.05) in chronic stressed animals compared to acute correlations were mapped. Blue lines indicate strengthened correlations, and red lines indicate disrupted connections, with line thickness representing strength of *z-*diff.

To identify potential network connections altered by continued exposure to restraint stress, Spearman correlation *r-*values underwent Fisher’s r-to-z transformation to compare correlation strength between groups. A *z*-difference (*z*-diff) for each inter-regional comparison was calculated ((*z*-diff of Chronic – *z*-diff of Acute) /standard error) and a two-tailed z-test was performed. Only connections with a *p*-value < 0.05 were graphed (**Fig. 9D**). In males, network-connectivity analysis demonstrated a disruption of connectivity between the BMA and LHb, the VTA and LC, and the Arc and LHA (**Fig. 9D, left**). In females, only the connections between the CeA and PVT were disrupted (**Fig. 9D, right**). While various network connections were strengthened over the course of chronic stress exposure between both males and females, the only overlapping connection were between the MeA and NTS (*z*-diff = 2.73 males, *z*-diff = 2.37 females). When compared to disruptions and enhanced connections following acute stress compared to controls (Supplemental Fig. 1), males suffered a disruption of connections between the BLA and PVT (Supplementary Fig. 1A) that is strengthened following chronic stress (**Fig. 9D left**). In females, of the 11 connections disrupted by acute stress compared to control (Supplementary Fig. 1B), 7 of those same connections are strengthened between exposure to chronic stress compared to acute (**Fig. 9D right**).

## Discussion

The results from the present study demonstrated that acute restraint stress caused broad neural activation in the brain, and that chronic (one hour daily for 10 days) restraint stress generally resulted in attenuated neural activity in a subset of those same stress-sensitive brain regions in both male and female mice. However, this expression of c-Fos-positive cells demonstrated a sex-dependent pattern in some regions of the hypothalamus, thalamus, midbrain, and hindbrain. The sex-associated difference in response to chronic versus acute stress was also seen via differences in attenuation of corticosterone between acute and chronic restraint groups. Together, these results support that male and female mice have both shared and divergent neural responses to acute and chronic restraint stress. Thus, chronic stress-induced desensitization, or lack-thereof, of neural activity in specific brain regions may contribute to sex differences in the development and prevalence of neuropsychiatric disorders^27–30^ and other stress-related (systemic) cardiovascular^23, 24, 47^ and gut inflammation diseases^25, 48^.

### Propensity for neural desensitization to stress is sex-specific in some brain regions

Our results suggest that brain regions where stress-induced neural activity fully desensitized following chronic restraint stress in males and females are primarily related to emotional processing of fear, anxiety, depression, and/or aversion including the LS^49, 50^, Arc^51–53^, BMA^54, 55^, MeA^56, 57^, and LHb^58, 59^. This normalization of emotional and behavioral control centers following chronic restraint stress aligns with the hypothesis that repeated *predictable* stress leads to habituation to the stressor^60^, decreasing the emotional component of stress responses. Interestingly, decreased neural activity, compared to control, of the LS, Arc, and MeA because of less predictable and/or more severe stressors, is associated with increased depression-like responses^53, 61, 62^. In contrast, a prolonged increase of LHb activity is associated with depression-like behaviors^58, 59, 63^. Our results demonstrated that females had a weaker increase in c-Fos expression from control compared to males in the BMA, MeA and LHb suggesting that restraint stress may be less emotionally stressful in females compared to males. Indeed, prior studies of chronic restraint stress have shown that males and females differ in severity of responses in measures of anhedonia^64^ and anxiety-like behaviors^65^, however a meta-analysis in 2022 found only 2 out of 57 chronic restraint stress protocols intending to model behavioral outcomes used female subjects^66^, limiting our general conclusions.

Simultaneously, there were several regions that did not fully desensitize neural reactivity to chronic restraint in both males and females. In these cases, c-Fos expression following chronic stress was increased compared to control mice but was decreased compared to their acutely stressed counterparts. These regions, the PVH and DMH, have key roles in the endocrine/HPA stress response^67–69^. In these regions we observed a decrease in neural activity in both males and females following chronic stress compared to acute stress and, in females, the magnitude of change in c-Fos expression following acute and chronic time points compared to control was smaller compared to males. Corticosterone responses, however, demonstrate an elevated level of corticosterone in females compared to males with no significant attenuation following chronic stress within sex (**Fig. 2**). This misaligned result could be explained by differences in the release of corticosterone related to PVH or DMH activation. Indeed, chemogenetic activation of PVH CRH neurons causes more corticosterone release in female mice than males^70^, additionally, a subset of DMH neurons play a role in inhibiting corticosterone release in response to emotional stressors^71^, thus the decrease in DMH activation in females compared to males could be an effect of decreased inhibition of the HPA axis. This, in tandem with the difference in corticosterone release by the PVH, could explain the observed sex-differences in corticosterone levels following restraint.

We found no desensitization of c-Fos expression in the LC (both males and females) and VTA (males only), with minimal desensitization in chronically stressed females that is not present when comparing the precent change of c-Fos expression (**Fig. 5C-D**). The VTA is an important region for reward and motivated behaviors that can be altered by stress^72, 73^, and is associated with stress-induced anxiety^74^. Restraint stress has been shown to increase firing of the VTA that is not attenuated over repeated exposure in males^75^ aligning with the lack of desensitization we observed. The LC is a key region for arousal^76–79^, via its norepinephrine (NE) projections to broad regions of the brain. PTSD and depression are characterized by symptoms of hyperarousal^80^, like insomnia, which is more pronounced in women with PTSD and depression^81, 82^. Surprisingly, no difference was found in the LC between males and females to acute stress or chronic stress in the present study, suggesting that sex differences may be altered by factors like ovarian hormones that regulate NE synthesis^83, 84^ and NE release in LC terminal regions^85–87^, rather than differences in LC neuronal activity.

### Main sex effects

While no regions were found to have increased c-Fos expression following chronic stress compared to acute stress, there were several regions where males and females differed in whether chronic stress-induced c-Fos was normalized or remained significantly increased compared to control expression. Of these regions, there was a main sex effect in either the raw c-Fos-positive count and/or the percent change of c-Fos expression in the LHA and BLA (**Fig. 8**). In both regions, the current study found that male c-Fos expression was only partially attenuated following chronic stress whereas female expression fully attenuated compared to same sex controls. The LHA is implicated in feeding, arousal, motivated behaviors, anxiety, and broadly both the development of and protection from depression-like phenotypes, depending on the activated subpopulation of neurons^88–97^. The BLA is also implicated in motivated behaviors, instrumental feeding and anxiety^98–103^, as well as conditioned fear learning, social behaviors, and defensive behaviors^102–104^, with respect to the specific subpopulation of neurons activated. Interestingly, the BLA and LHA have recently been proposed to integrate information together to bias motivated behaviors towards relevant stimuli^105^. While our correlational analysis did not demonstrate a significant correlation between these two regions, the parallel sex differences may suggest sex-specific behavioral outcomes related to stress-induced motivated behaviors^106^, feeding^107–109^, and anxiety^42, 65^, all of which require more investigation.

### Changes in region interconnectedness

When investigating the potential interconnectedness between the established regions, it is generally apparent that the female control mice had similar levels of c-Fos expression across brain regions resulting in many significant positive correlations, and even those that were not significant trended towards a positive correlation rather than a negative correlation. Whereas males had a handful of significant positive correlations and ∼50 % of correlations trending positive and ∼50 % negative. While the overall regions we quantified differed from those studied in a similar study^110^, our findings in shared regions seem to align with overall opposite connections in male and female regional interconnectedness in the control group, and a divergence from control correlations during acute stress^110^. During acute stress, males had an increase of some of the correlations, becoming significantly correlated (both positive and negative), while other connections, that had been trending towards positive or negative correlations during control conditions, switched, and became significantly correlated in the oppositive direction. However, in females, a few of the positive correlations are conserved, but many of the positive correlations, and possible interconnectedness, are disrupted. Only the correlations between the PVT and LHb are increased by acute stress. However, based on c-Fos expression, we know these regions aren’t being inactivated, but previous studies have shown that stress protocols inducing depression cause a shift from activating positive valance sites (like the VTA and NAc) to regions associated with negative valence (like the CeA and LHb)^111–114^. Following chronic stress, males continue to increase the number of significant positive correlations as well as overall positive correlations. Females also increase the number of positive correlations compared to acute stress but do not regain the level of positive correlations seen in control mice. Without investigating the biological significance of every correlation and mapping each correlation change between all three conditions, we still see an overall trend demonstrating brain wide changes in interconnectedness that are likely involved in the known behavioral and physiological outcomes of restraint stress. The divergent pattern in male versus female mice is apparent with only a few correlations being shared between them. Namely, in controls 3 out of the 5 male positive correlations are also found in females. During acute stress, the positive correlation between the CeA and LS is found in both sexes, and during chronic stress, only the positive correlation between the PVT and LS is shared. Interestingly, the CeA and LS have both been shown to be downstream targets of the infralimbic cortex (IL), modulating both anxiolytic and anxiogenic behaviors, respectively^115^. This may demonstrate a shared function in male and female brains where the IL is responding to the stressful environment and enacting defensive behaviors, like freezing, while also exerting top-down control of anxiety via the CeA. This connection is lost in females during chronic stress suggesting that the balanced activation of both regions is lost, potentially resulting in aberrant responses like depression.

While the interconnectedness of regions following each condition is an interesting avenue of investigation for the bio-behavioral output at those time points, how these network connections change from acute to chronic stress remains elusive. We show here that in males, the interconnectedness of 8 paired regions is strengthened by the end of chronic restraint stress and 3 pairs were disrupted compared to following a single restraint. Of these regions that are now more strongly interconnected following chronic stress, 3 of the pairs (the BLA and MeA, BLA and PVT, and MeA and CeA) had a positive correlation (not significant) at baseline suggesting that the connectivity of these regions may have strengthened because of desensitization of one or more regions to stress over time. However, as these regions are all highly associated with fear, anxiety, and depression, strengthened connections may be a maladaptive outcome^56, 116^. Additionally, of the three paired regions found to be disrupted over extended stress exposure, only the Arc and LHA connection had originally been a negative correlation, suggesting another network returning to similar level of connectedness as seen at baseline. In females, 10 paired regions strengthened connections over chronic stress and one pair (CeA and PVT) was disrupted. Of these strengthened pairs, all 10 were positively associated with each other (5/10 statistically significant), suggesting that most of these regions were desensitized and the interconnectedness was normalized by the end of chronic stress. The CeA and PVT, however, were positively ^117, 118^ correlated at baseline and their interconnectedness was disrupted by chronic stress. The connectivity of the CeA and PVT are implicated in stress-induced depression and insomnia-related symptoms^117, 118^, potentially supporting increased depressive-like symptoms following stress in females despite the significant recovery of many neural connections. Interestingly, while we seem to see more desensitization and normalization of network connectivity in females than males, we still see a heightened level of corticosterone in females compared to males emphasizing the complexity of long-term stress exposure. These results further demonstrate that the desensitization of neural responses may differ between males and females^19, 20^. While some neural circuits may return to similar, or enhanced, levels of connectivity, the continued disruption of select connections and the differences between sexes warrant further exploration for their role in chronic stress-induced behavioral and physiological outcomes.

## Conclusion

In the present study, we found broad brain regions in both male and female mice that were activated by acute restraint stress, and in most of these brain regions, the degree of activation was at least partially attenuated after chronic restraint stress, suggesting a desensitization to the stressor. Sex differences of c-Fos expression to acute stress and chronic stress were found in some hypothalamus, amygdala and midbrain regions. An indirect measure of regional connectedness also showed distinctly different patterns of correlation between sexes that suggest many potentially interesting circuits that may be differentially activated in males and females over acute to chronic restraint stress exposure. Together this study demonstrates that male and female mice respond and/or adapt to acute and chronic stress differently, via neuronal pathways in the amygdala, hypothalamus, midbrain, and hindbrain, and that such changes possibly contribute to the sex-dependent prevalence of neuropsychiatric disorders. It also underscores the importance of including both male and female mice in stress protocols, as there are few studies alluding to the potential behavioral and physiological outcomes of neural differences.

---

**Supplemental Figure 1.**
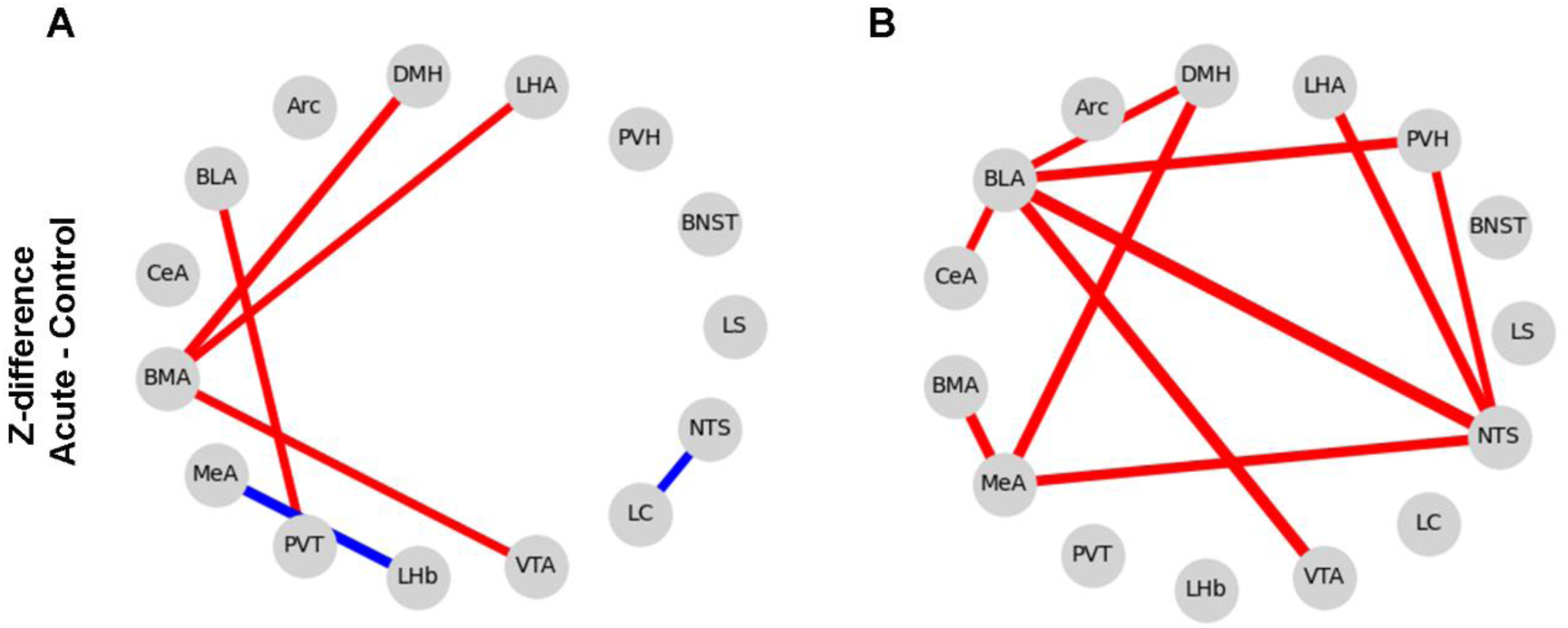
Change in regional interconnectedness between acute and control mice. The strongest changes in network connectivity (as indicated by the difference in Fisher’s transformed *z*-scores, i.e. *z-*diff, p < 0.05) in acute stressed males **(A)** and females **(B)** compared to control correlations were mapped. Blue lines indicate strengthened correlations, and red lines indicate disrupted connections, with line thickness representing strength of *z-*diff.

**Supplemental Table 1.** Effects of sex and stress on c-Fos-positive cells per region. Two-way ANOVA for main effects of the interaction between sex and stress, sex, and stress are reported as *F-* and *p-*values. For each region, subsequent post hoc Tukey’s multiple comparisons for comparisons of stress groups within sex and between sex are reported as *p*-values. Significant *p*-values (p < 0.05) are bolded. Any post hoc comparisons statistically identified as significant but without a significant main effect are in red and not reported in related graphs.

|  | Two-Way ANOVA |  |  | Tukey's Multiple Comparisons Test |  |  |  |  |
| --- | --- | --- | --- | --- | --- | --- | --- | --- |
|  | Comparison | F value | p value | Within Sex |  |  | Between Sex |  |
|  |  |  |  |  | Male | Female |  |  |
| LS | Interaction | F (2, 42) = 0.2975 | *p*=0.7442 | Ctrl vs Acute | ***p*<0.0001** | ***p*<0.0001** | Ctrl | *p*=0.9984 |
|  | Sex | F (1, 42) = 0.6150 | *p*=0.4373 | Ctrl vs Chronic | *p*=0.3895 | *p*=0.5684 | Acute | *p*=0.2963 |
|  | Stress | F (2, 42) = 39.86 | ***p*<0.0001** | Acute vs Chronic | ***p*<0.0001** | ***p*=0.0002** | Chronic | *p*=0.7636 |
| BNST | Interaction | F (2, 42) = 1.027 | *p*=0.3669 | Ctrl vs Acute | ***p*<0.0001** | ***p*<0.0001** | Ctrl | *p*=0.6983 |
|  | Sex | F (1, 42) = 0.03318 | *p*=0.8563 | Ctrl vs Chronic | *p*=0.1023 | ***p*=0.0295** | Acute | *p*=0.3135 |
|  | Stress | F (2, 42) = 37.35 | ***p*<0.0001** | Acute vs Chronic | ***p*<0.0001** | ***p*=0.026** | Chronic | *p*=0.3498 |
| PVN | Interaction | F (2, 42) = 0.7323 | *p*=0.4868 | Ctrl vs Acute | ***p*<0.0001** | ***p*<0.0001** | Ctrl | *p*=0.8823 |
|  | Sex | F (1, 42) = 0.1470 | *p*=0.7034 | Ctrl vs Chronic | ***p*=0.0007** | ***p*=0.0114** | Acute | *p*=0.5544 |
|  | Stress | F (2, 42) = 86.21 | ***p*<0.0001** | Acute vs Chronic | ***p*<0.0001** | ***p*<0.0001** | Chronic | *p*=0.2729 |
| LHA | Interaction | F (2, 42) = 2.863 | *p*=0.0683 | Ctrl vs Acute | ***p*<0.0001** | ***p*<0.0001** | Ctrl | *p*=0.4529 |
|  | Sex | F (1, 42) = 0.005804 | *p*=0.9396 | Ctrl vs Chronic | ***p*=0.0005** | *p*=0.3653 | Acute | *p*=0.2784 |
|  | Stress | F (2, 42) = 59.71 | ***p*<0.0001** | Acute vs Chronic | ***p*=0.0049** | ***p*<0.0001** | Chronic | *p*=0.0534 |
| DMH | Interaction | F (2, 42) = 0.1096 | *p*=0.8964 | Ctrl vs Acute | ***p*<0.0001** | ***p*<0.0001** | Ctrl | *p*=0.3242 |
|  | Sex | F (1, 42) = 4.914 | ***p*=0.0321** | Ctrl vs Chronic | ***p*=0.0002** | ***p*<0.0001** | Acute | *p*=0.1076 |
|  | Stress | F (2, 42) = 112.8 | ***p*<0.0001** | Acute vs Chronic | ***p*<0.0001** | ***p*<0.0001** | Chronic | *p*=0.2378 |
| Arc | Interaction | F (2, 42) = 0.7215 | *p*=0.4920 | Ctrl vs Acute | ***p*<0.0001** | ***p*<0.0001** | Ctrl | *p*=0.4113 |
|  | Sex | F (1, 42) = 0.1872 | *p*=0.6675 | Ctrl vs Chronic | *p*=0.1118 | *p*=0.0761 | Acute | *p*=0.4724 |
|  | Stress | F (2, 42) = 96.85 | ***p*<0.0001** | Acute vs Chronic | ***p*<0.0001** | ***p*<0.0001** | Chronic | *p*=0.5227 |
| BLA | Interaction | F (2, 42) = 3.428 | ***p*=0.0418** | Ctrl vs Acute | ***p*<0.0001** | ***p*<0.0001** | Ctrl | *p*=0.1071 |
|  | Sex | F (1, 42) = 0.2092 | *p*=0.6497 | Ctrl vs Chronic | ***p*=0.008** | *p*=0.9439 | Acute | *p*=0.3302 |
|  | Stress | F (2, 42) = 52.02 | ***p*<0.0001** | Acute vs Chronic | ***p*=0.0007** | ***p*<0.0001** | Chronic | *p*=0.0729 |
| CeA | Interaction | F (2, 42) = 0.7521 | *p*=0.4776 | Ctrl vs Acute | ***p*<0.0001** | ***p*<0.0001** | Ctrl | *p*=0.5931 |
|  | Sex | F (1, 42) = 0.02263 | *p*=0.8812 | Ctrl vs Chronic | *p*=0.1771 | *p*=0.9783 | Acute | *p*=0.7825 |
|  | Stress | F (2, 42) = 37.46 | ***p*<0.0001** | Acute vs Chronic | ***p*=0.0006** | ***p*<0.0001** | Chronic | *p*=0.2877 |
| BMA | Interaction | F (2, 42) = 0.7791 | *p*=0.4653 | Ctrl vs Acute | ***p*<0.0001** | ***p*<0.0001** | Ctrl | *p*=0.1429 |
|  | Sex | F (1, 42) = 1.746 | *p*=0.1935 | Ctrl vs Chronic | *p*=0.6064 | *p*=0.7341 | Acute | *p*=0.3166 |
|  | Stress | F (2, 42) = 57.46 | ***p*<0.0001** | Acute vs Chronic | ***p*<0.0001** | ***p*<0.0001** | Chronic | *p*=0.8285 |
| MeA | Interaction | F (2, 42) = 0.9849 | *p*=0.3820 | Ctrl vs Acute | ***p*<0.0001** | ***p*<0.0001** | Ctrl | *p*=0.1999 |
|  | Sex | F (1, 42) = 2.401 | *p*=0.1288 | Ctrl vs Chronic | *p*=0.6069 | *p*=0.8311 | Acute | *p*=0.1131 |
|  | Stress | F (2, 42) = 129.7 | ***p*<0.0001** | Acute vs Chronic | ***p*<0.0001** | ***p*<0.0001** | Chronic | *p*=0.814 |
| PVT | Interaction | F (2, 42) = 1.611 | *p*=0.2117 | Ctrl vs Acute | ***p*<0.0001** | ***p*=0.0002** | Ctrl | *p*=0.8095 |
|  | Sex | F (1, 42) = 6.160 | ***p*=0.0172** | Ctrl vs Chronic | ***p*=0.0018** | ***p*=0.0296** | Acute | ***p*=0.0083** |
|  | Stress | F (2, 42) = 32.84 | ***p*<0.0001** | Acute vs Chronic | ***p*=0.0057** | *p*=0.1812 | Chronic | *p*=0.205 |
| LHb | Interaction | F (2, 42) = 0.2248 | *p*=0.7997 | Ctrl vs Acute | ***p*<0.0001** | ***p*<0.0001** | Ctrl | *p*=0.6322 |
|  | Sex | F (1, 42) = 0.1845 | *p*=0.6697 | Ctrl vs Chronic | *p*=0.0835 | *p*=0.0707 | Acute | *p*=0.7675 |
|  | Stress | F (2, 42) = 113.5 | ***p*<0.0001** | Acute vs Chronic | ***p*<0.0001** | ***p*<0.0001** | Chronic | *p*=0.5788 |
| VTA | Interaction | F (2, 42) = 2.029 | *p*=0.1441 | Ctrl vs Acute | ***p*<0.0001** | ***p*<0.0001** | Ctrl | *p*=0.604 |
|  | Sex | F (1, 42) = 1.737 | *p*=0.1947 | Ctrl vs Chronic | ***p*<0.0001** | ***p*<0.0001** | Acute | ***p*=0.0271** |
|  | Stress | F (2, 42) = 75.33 | ***p*<0.0001** | Acute vs Chronic | *p*=0.9956 | ***p*=0.0245** | Chronic | *p*=0.5993 |
| LC | Interaction | F (2, 42) = 0.1151 | *p*=0.8916 | Ctrl vs Acute | ***p*<0.0001** | ***p*<0.0001** | Ctrl | *p*=0.939 |
|  | Sex | F (1, 42) = 0.4788 | *p*=0.4928 | Ctrl vs Chronic | ***p*<0.0001** | ***p*<0.0001** | Acute | *p*=0.4555 |
|  | Stress | F (2, 42) = 76.12 | ***p*<0.0001** | Acute vs Chronic | *p*=0.9329 | *p*=0.9995 | Chronic | *p*=0.7146 |
| NTS | Interaction | F (2, 41) = 0.5151 | *p*=0.6012 | Ctrl vs Acute | ***p*<0.0001** | ***p*<0.0001** | Ctrl | *p*=0.7849 |
|  | Sex | F (1, 41) = 0.06604 | *p*=0.7985 | Ctrl vs Chronic | *p*=0.303 | ***p*=0.0232** | Acute | *p*=0.8042 |
|  | Stress | F (2, 41) = 64.09 | ***p*<0.0001** | Acute vs Chronic | ***p*<0.0001** | ***p*<0.0001** | Chronic | *p*=0.3312 |

**Supplemental Table 2.**
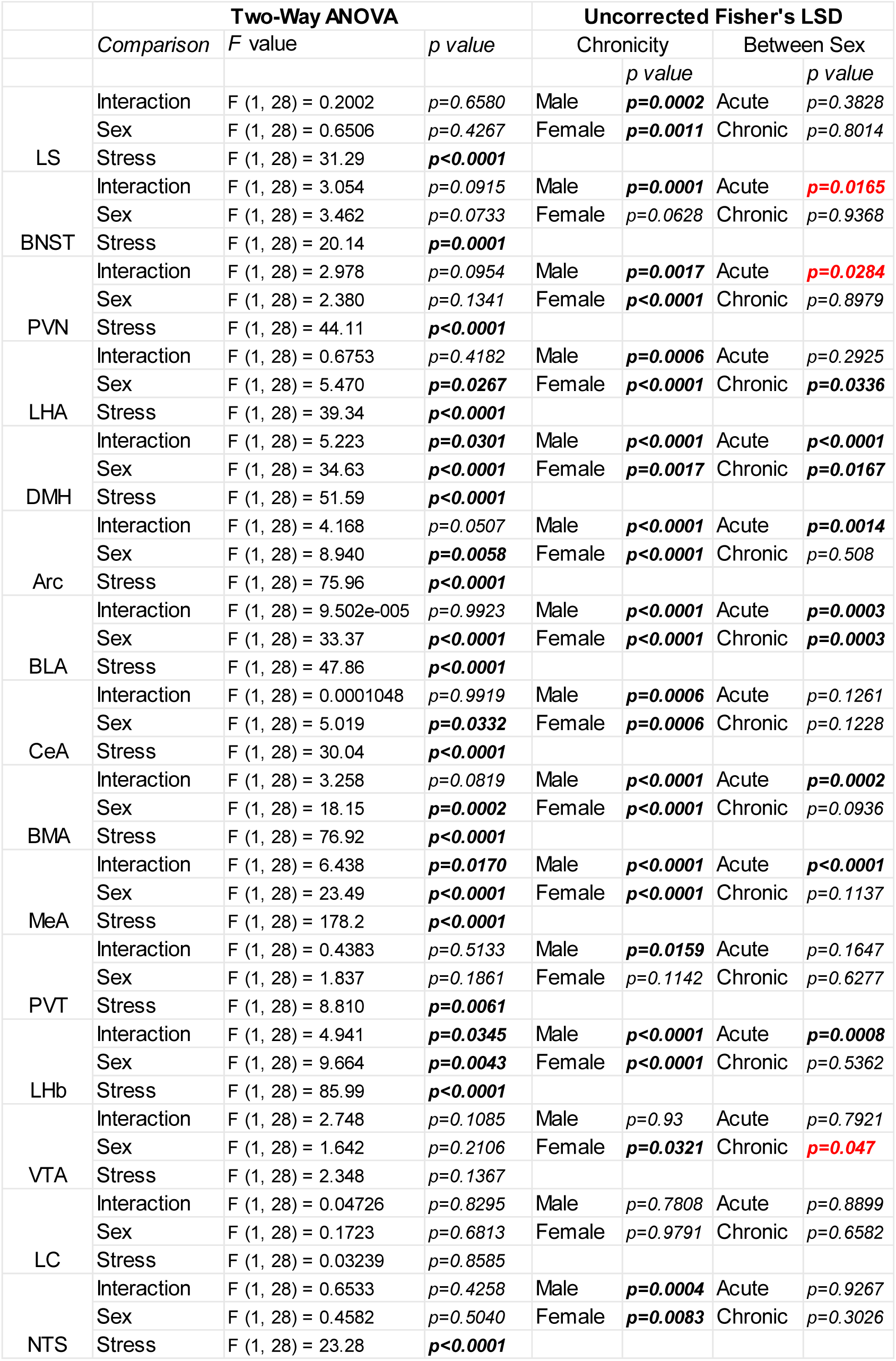
Effects of sex and stress on percent change of c-Fos-positive expression per region. Two-way ANOVA for main effects of the interaction between sex and stress, sex, and stress are reported as *F-* and *p-*values. For each region, subsequent post hoc Uncorrected Fisher’s LSD for comparisons of acute versus chronic stress per sex and between are reported as *p*-values. Significant *p*-values (p < 0.05) are bolded. Any post hoc comparisons statistically identified as significant but without a significant main effect are in red and not reported in related graphs.

|  | Two-Way ANOVA |  |  | Uncorrected Fisher's LSD |  |  |  |
| --- | --- | --- | --- | --- | --- | --- | --- |
|  | Comparison | F value | p value | Chronicity |  | Between Sex |  |
|  |  |  |  |  | p value |  | p value |
| LS | Interaction | F (1, 28) = 0.2002 | $p=0.6580$ | Male | $p=0.0002$ | Acute | $p=0.3828$ |
| | Sex | F (1, 28) = 0.6506 | $p=0.4267$ | Female | $p=0.0011$ | Chronic | $p=0.8014$ |
| | Stress | F (1, 28) = 31.29 | $p<0.0001$ | | | | |
| BNST | Interaction | F (1, 28) = 3.054 | $p=0.0915$ | Male | $p=0.0001$ | Acute | $p=0.0165$ |
| | Sex | F (1, 28) = 3.462 | $p=0.0733$ | Female | $p=0.0628$ | Chronic | $p=0.9368$ |
| | Stress | F (1, 28) = 20.14 | $p=0.0001$ | | | | |
| PVN | Interaction | F (1, 28) = 2.978 | $p=0.0954$ | Male | $p=0.0017$ | Acute | $p=0.0284$ |
| | Sex | F (1, 28) = 2.380 | $p=0.1341$ | Female | $p<0.0001$ | Chronic | $p=0.8979$ |
| | Stress | F (1, 28) = 44.11 | $p<0.0001$ | | | | |
| LHA | Interaction | F (1, 28) = 0.6753 | $p=0.4182$ | Male | $p=0.0006$ | Acute | $p=0.2925$ |
| | Sex | F (1, 28) = 5.470 | $p=0.0267$ | Female | $p<0.0001$ | Chronic | $p=0.0336$ |
| | Stress | F (1, 28) = 39.34 | $p<0.0001$ | | | | |
| DMH | Interaction | F (1, 28) = 5.223 | $p=0.0301$ | Male | $p<0.0001$ | Acute | $p<0.0001$ |
| | Sex | F (1, 28) = 34.63 | $p<0.0001$ | Female | $p=0.0017$ | Chronic | $p=0.0167$ |
| | Stress | F (1, 28) = 51.59 | $p<0.0001$ | | | | |
| Arc | Interaction | F (1, 28) = 4.168 | $p=0.0507$ | Male | $p<0.0001$ | Acute | $p=0.0014$ |
| | Sex | F (1, 28) = 8.940 | $p=0.0058$ | Female | $p<0.0001$ | Chronic | $p=0.508$ |
| | Stress | F (1, 28) = 75.96 | $p<0.0001$ | | | | |
| BLA | Interaction | F (1, 28) = 9.502e-005 | $p=0.9923$ | Male | $p<0.0001$ | Acute | $p=0.0003$ |
| | Sex | F (1, 28) = 33.37 | $p<0.0001$ | Female | $p<0.0001$ | Chronic | $p=0.0003$ |
| | Stress | F (1, 28) = 47.86 | $p<0.0001$ | | | | |
| CeA | Interaction | F (1, 28) = 0.0001048 | $p=0.9919$ | Male | $p=0.0006$ | Acute | $p=0.1261$ |
| | Sex | F (1, 28) = 5.019 | $p=0.0332$ | Female | $p=0.0006$ | Chronic | $p=0.1228$ |
| | Stress | F (1, 28) = 30.04 | $p<0.0001$ | | | | |
| BMA | Interaction | F (1, 28) = 3.258 | $p=0.0819$ | Male | $p<0.0001$ | Acute | $p=0.0002$ |
| | Sex | F (1, 28) = 18.15 | $p=0.0002$ | Female | $p<0.0001$ | Chronic | $p=0.0936$ |
| | Stress | F (1, 28) = 76.92 | $p<0.0001$ | | | | |
| MeA | Interaction | F (1, 28) = 6.438 | $p=0.0170$ | Male | $p<0.0001$ | Acute | $p<0.0001$ |
| | Sex | F (1, 28) = 23.49 | $p<0.0001$ | Female | $p<0.0001$ | Chronic | $p=0.1137$ |
| | Stress | F (1, 28) = 178.2 | $p<0.0001$ | | | | |
| PVT | Interaction | F (1, 28) = 0.4383 | $p=0.5133$ | Male | $p=0.0159$ | Acute | $p=0.1647$ |
| | Sex | F (1, 28) = 1.837 | $p=0.1861$ | Female | $p=0.1142$ | Chronic | $p=0.6277$ |
| | Stress | F (1, 28) = 8.810 | $p=0.0061$ | | | | |
| LHb | Interaction | F (1, 28) = 4.941 | $p=0.0345$ | Male | $p<0.0001$ | Acute | $p=0.0008$ |
| | Sex | F (1, 28) = 9.664 | $p=0.0043$ | Female | $p<0.0001$ | Chronic | $p=0.5362$ |
| | Stress | F (1, 28) = 85.99 | $p<0.0001$ | | | | |
| VTA | Interaction | F (1, 28) = 2.748 | $p=0.1085$ | Male | $p=0.93$ | Acute | $p=0.7921$ |
| | Sex | F (1, 28) = 1.642 | $p=0.2106$ | Female | $p=0.0321$ | Chronic | $p=0.047$ |
| | Stress | F (1, 28) = 2.348 | $p=0.1367$ | | | | |
| LC | Interaction | F (1, 28) = 0.04726 | $p=0.8295$ | Male | $p=0.7808$ | Acute | $p=0.8899$ |
| | Sex | F (1, 28) = 0.1723 | $p=0.6813$ | Female | $p=0.9791$ | Chronic | $p=0.6582$ |
| | Stress | F (1, 28) = 0.03239 | $p=0.8585$ | | | | |
| NTS | Interaction | F (1, 28) = 0.6533 | $p=0.4258$ | Male | $p=0.0004$ | Acute | $p=0.9267$ |
| | Sex | F (1, 28) = 0.4582 | $p=0.5040$ | Female | $p=0.0083$ | Chronic | $p=0.3026$ |
| | Stress | F (1, 28) = 23.28 | $p<0.0001$ | | | | |

